# AAV Capsid Chimeras with Enhanced Infectivity reveal a core element in the AAV Genome critical for both Cell Transduction and Capsid Assembly

**DOI:** 10.1101/2020.10.13.337261

**Authors:** Lydia Viney, Tilmann Bürckstümmer, Mario Mietzsch, Modassir Choudhry, Tom Henley, Mavis Agbandje-McKenna

## Abstract

Adeno-associated viruses (AAV) have attracted significant attention in the field of gene and cell therapy due to highly effective delivery of therapeutic genes into human cells. The ability to generate recombinant AAV vectors compromised of unique or substituted protein sequences has led to the development of capsid variants with improved therapeutic properties. Seeking a novel AAV capable of enhanced transduction of human T cells for applications in immunotherapy, we have developed a unique capsid variant termed AAV *X-Vivo* (AAV-XV) that is a chimera of AAV12 VP1/2 sequences and the VP3 sequence of AAV6. This AAV chimera showed enhanced infection of human primary T cells and hematopoietic stem cells, and superiority over wildtype AAV6 for the genomic integration of DNA sequences either by AAV alone or in combination with CRISPR gene editing. AAV-XV demonstrated transduction efficiency equivalent to AAV6 at multiplicities of infection 2 logs lower, enabling T cell engineering at low AAV doses. Analyzing the protein coding sequence of AAV-XV revealed disruptions within the assembly-activating protein (AAP) which likely accounted for observed lower virus yield. A series of genome alterations reverting the AAP sequence back to wildtype had a negative impact on the enhanced transduction seen with AAV-VX, indicating overlapping functions within this sequence for both viral assembly and effective T cell transduction. Our findings show that AAV-XV is highly efficient at T cell engineering at low AAV dose and demonstrates the importance of AAP coding region in both viral particle assembly and cell infection.

**IMPORTANCE:** A major hurdle to the therapeutic potential of AAV in gene therapy lies in achieving clinically meaningful AAV doses, and secondarily, ability to manufacture commercially viable titers of AAV to support this. By virtue of neutralizing antibodies against AAV that impede patient repeat-dosing, the dose of AAV for *in vivo* gene delivery has been high, which has resulted in unfortunate recent safety concerns and deaths in patients given higher-dose AAV gene therapy. We have generated a new AAV variant possessing a unique combination of capsid proteins for ex-vivo application termed AAV-XV, which delivers high levels of cell transduction and gene delivery at a lower MOI. Furthermore, we demonstrate a novel finding, and an important consideration for recombinant AAV design, that a region of the AAV genome encoding the capsid protein and AAP gene is critical for both virus yield and the enhancement of infection/transduction.

## INTRODUCTION

The Adeno-associated viruses (AAVs) are one of the most widely developed and actively studied vehicles for gene and cell therapy, and have shown remarkable promise in numerous clinical trials for multiple human disorders (1). As a vector for gene-delivery into human cells, these single-stranded DNA viruses show broad tropism, with multiple serotypes identified for transducing cells from many different tissue types (1). Their non-pathogenicity coupled with long-term transgene expression, makes AAV attractive as a therapeutic technology for *in vivo* gene therapy (2). Despite broadly low innate immunogenicity, concerns over humoral immune responses against AAV capsids, observed in recent clinical trials, have been raised, particularly associated with high vector doses (3, 4). This *in vivo* limitation, as well as the high doses or multiplicities of infection (MOI) of virus required for sufficient cell transduction, and the need to expand the repertoire of transducible tissue types addressable with AAV, is motivating further development of recombinant AAV technology.

There are 13 naturally occurring AAV serotypes and numerous AAV isolates (5), each with unique capsid viral protein (VP) sequences and transduction profiles in different tissues (6). As an example, AAV6 is consistently better than other serotypes in *ex vivo* transduction of human immune cells (7, 8). Each serotypes’ distinct VP sequences assemble in strict T=1 icosahedral arrangement that enables packaging of the AAV genome into an infectious virion (9). Novel variants of AAV are also being identified from sequencing experiments in different cell types, such as within CD34+ hematopoietic stem cells (10). Furthermore, uniquely engineered AAV vectors with enhanced transduction efficiencies have been developed (11) through capsid mutation by rational design (12, 13), directed evolution (14), or by combining different serotypes through capsid shuffling (15). Thus, combining sequences from divergent serotypes, or specific mutations of surface exposed capsid residues known to facilitate viral entry into cells, may be an effective route to improve the infectious properties of AAV.

While AAV vector transduction can lead to high levels of transient transgene expression by episomal genomes, integration into the host genome typically occurs at very low frequency (16). The stable genomic integration of AAV donor vectors can be increased significantly via the combination of AAV vectors with CRISPR/Cas9 gene editing (17). A targeted double-strand break (DSB) introduced by Cas9 at a specific location within the genome can be effectively repaired with an AAV template designed with homology to the target locus, via the pathway of homology-directed repair (HDR) (17). This AAV + CRISPR combination approach has been effectively used by us and others to perform genetic engineering of difficult to target cell types such as primary human T cells at levels of efficiency that are therapeutically relevant (18). However, despite the advances in AAV vector engineering, capsid evolution, and use of synergistic technologies such as CRISPR, high AAV dose MOIs, typically 1×10^6^ virus particles per cell, are still required for gene delivery into cells being modified for either research or therapeutic applications (19). This makes AAV a costly technology to deploy at scale for cell therapies, and as mentioned, may preclude *in vivo* efficacy due to potentially toxic high doses required for gene therapy.

To address the above limitations, a unique AAV capsid variant with enhanced transduction of human T cells was developed to improve the efficiency of *ex vivo* gene delivery. A series of capsid variants, were engineered, via rational design, by substituting the VP1 unique (VP1u) and VP1/2-common region sequences of AAV6 with those from divergent AAV serotypes such as AAV4, AAV5, AAV11, and AAV12 to create chimeric AAV6 vectors. Analysis of the resulting chimeras, for performance in transduction assays using primary human T cells, found several variants that achieved levels of transduction 100-fold higher than wild-type AAV6 at similar MOI or 10- to 30-fold higher at 2 log lower doses. The best performing variant was named AAV *X-Vivo* (AAV-XV). For this variant, an overlapping region of the *cap* open reading frame (ORF) encoding VP2 and the Assembly-Activating Protein (AAP), was observed to be critical for both optimal vector yield and efficient cellular infectivity. Our data demonstrates that AAV-XV is highly efficient at cellular transduction and a broadly relevant recombinant vector for the *ex-vivo* engineering of human T cells for immunotherapy and has potential as an efficient AAV for *in vivo* gene delivery at low doses.

## RESULTS

### Generation of new chimeric AAV6 capsid variants

To engineer capsid variants with the potential to transduce human T cells at high efficiencies, 7 chimeric AAV capsid sequences, for which AAV6 provided the VP3, were generated (Supplementary Fig. S1). Each variant incorporated sequence from the VP1u or VP1u+VP1/2 common regions of AAV4, AAV5, or AAV12 or VP1u+VP1/2 common region of AAV11. The serotypes substituted represent the most diverse based on the pairwise comparison of the VP1u and VP1/2 common regions from AAV1 through AAV13 with AAV6 (Supplementary Fig. S1), with the intent to create maximum diversity in the resulting chimeras. A multiple sequence alignment between the selected serotypes and AAV6 showed conservation of functional regions including a PLA2 motif (20), a calcium binding motif (21), and basic residue clusters that serve as nuclear localization sequences (NLS) (22), although positioning of the latter was different for AAV5 (Fig. 1A). This alignment showed a higher sequence identity for the VP1u region compared to the VP1/2 common region (Supplementary Fig. S1). Recombinant AAV vectors utilizing wild-type AAV6 and the 7 variant capsids (Fig. 1B), packaging a donor template with a NanoLuc luciferase gene, were produced and evaluated for their performance as *ex vivo* gene-delivery vectors in human T and stem cells.

**Figure 1.**
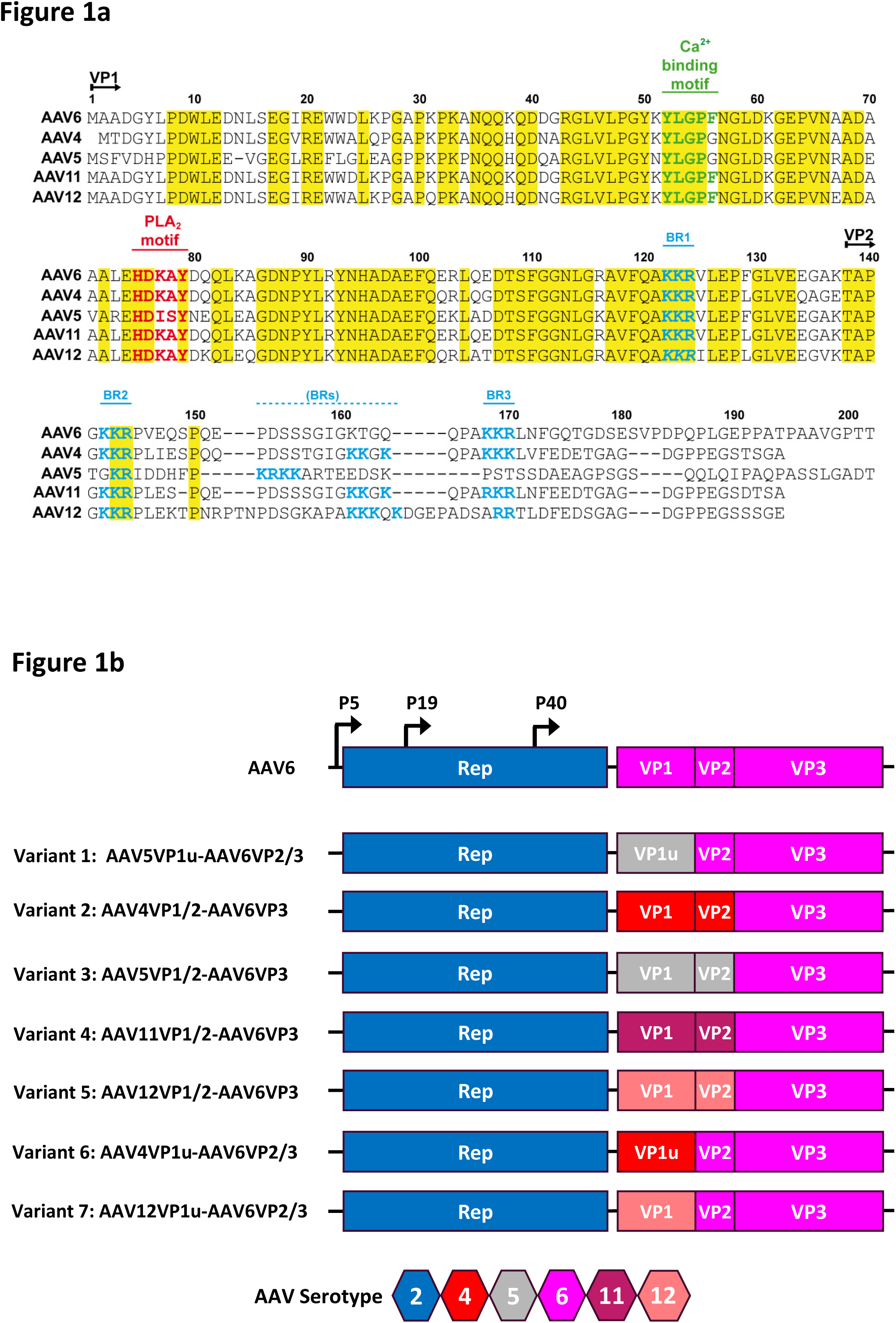
Design of recombinant AAV6 capsid variants. (A) Alignment of the VP1u and VP1/2 common region sequences of AAV4, AAV5, AAV6, AAV11, and AAV12 showing conservation of functionally important motifs. (B) Schematic diagram of the AAV2 Rep and the AAV serotypes contributing VP1, VP2, and VP3 to the AAV6 variants. The approximate positions of promoters are indicated. A serotype color key is provided.

### Chimeric AAV6 capsid variants outperform wildtype as donors for CRISPR genomic integration in human T and stem cells at low MOI

To assess the impact of the VP sequence substitutions into AAV6 on infectivity and genomic integration efficiencies, CD3+ T cells were electroporated with Cas9 mRNA and a sgRNA targeting the AAVS1 locus, followed by infection with the rAAV chimeras (Fig. 1B). The rAAV vectors delivered a NanoLuc expression cassette flanked by homology arms allowing targeted integration at the AAVS1 locus (23) (Fig. 2A). An MOI titration of these variants monitored 7- and 14-days post infection and CRISPR transfection showed 3 of the 7 chimeras (variants 2, 4, and 5), with significantly enhancement transduction and genomic integration of the targeting construct (1.5 to 2 logs) compared to wild-type AAV6 (Fig. 2B). Interestingly, at a low MOI of 1×10^4^ genome copies per cell (gc/cell), the best variant, 5, named AAV-XV, achieved luciferase levels that required a 100-fold higher MOI of wild-type AAV6 vectors (1×10^6^). Variants 2, 4, and 5 possess the VP1u+VP1/2 common region of AAV4, and AAV11, and AAV12, respectively (Fig. 1B), and unlike AAV5 and AAV6, contain an additional basic cluster (Fig. 1A). In contrast, the variants containing only the VP1u region of AAV4, and AAV11, and AAV12 did not display enhanced transduction and genomic integration suggesting that their VP1/2 common region is the determinant of this phenotype (Fig. 1B and 2B). Notably, the variant with substituted VP1+VP1/2 common region of AAV5 (variant 3) did not enhance transduction and was arguably the worse performing vector (Fig. 2B).

**Figure 2.**
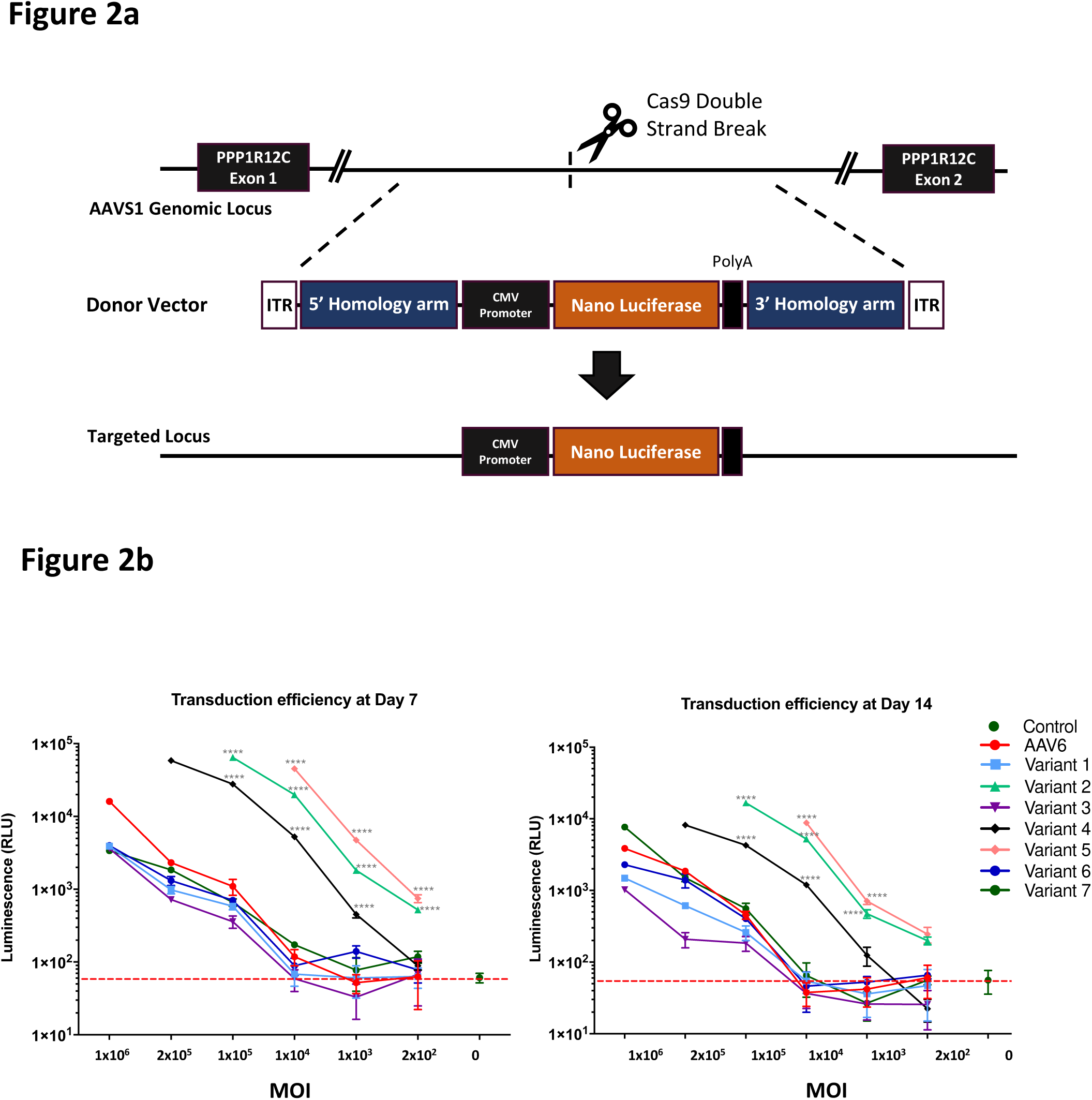
Transduction efficiency and CRISPR-mediated genomic integration of AAV6 capsid variants in human T cells. (A) Schematic of the AAV6 targeting vector for the integration of a luciferase expression cassette into the PPP1R12C (AAVS1) genomic locus. Diagram indicates the relative position of CRISPR-mediated genomic cleavage and the resulting modified locus upon homology-directed repair with the AAV6-luciferase vector. (B) MOI dose-titration of wild-type AAV6 and chimeric AAV6 capsid variants measuring CRISPR-mediated genomic integration of a luciferase reporter gene in human CD3+ T cells at 7 and 14 d post-infection. Statistical significance was determined by a one-way ANOVA test for multiple comparisons, ****P<0.0001.

Further analysis of the 3 best performing capsid variants in both human cytotoxic CD8+ T cells and CD34+ HSCs confirmed higher transduction efficiencies and luciferase targeting to the AAVS1 genomic site of up to 2 logs (Fig. 3A). This enhanced performance was also observed in the absence of a CRISPR-mediated DSB, with variant 5 showing a 1 log increase in transduction efficiency over AAV6 when measured up to 21 days post infection (Fig. 3B). Again, this variant achieved similar levels of T cell transduction at the lower MOI of 1×10^4^ gc/cell as AAV6 at MOI of 1×10^6^ gc/cell. AAV-XV thus demonstrated superior infectivity and transduction performance at low doses compared to the wild-type AAV6 sequence from which it was derived.

**Figure 3.**
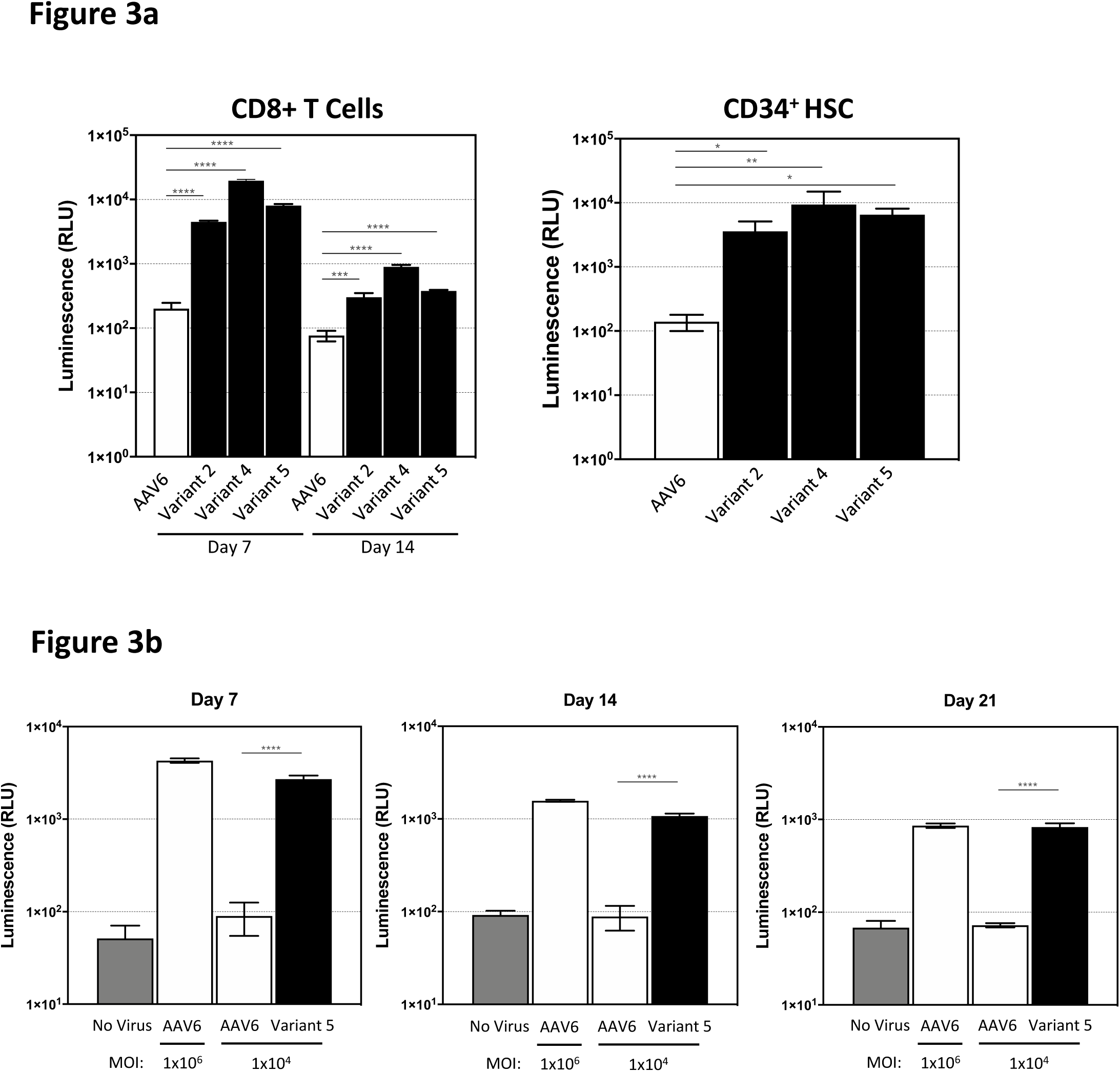
Low MOI transduction of human CD8+ T cells and HSCs by AAV6 capsid variants. (A) Comparison of CRISPR-mediated genomic integration by wild-type AAV6 and the top three performing AAV6 capsid variants at low MOI (1×10^4^ / cell) in primary CD8 T cells (left) and human CD34+ HSCs (right). (B) Comparison of transduction efficiency of capsid variant 5 (AAV12VP1/2-AAV6VP3) at low MOI (1×10^4^ / cell) and wild-type AAV6 at high MOI (1×10^6^ / cell) in CD8 T cells over a 3-week period in the absence of CRISPR gene editing. Statistical significance was determined by an unpaired T test, or one-way ANOVA test for multiple comparisons, *P<0.05, **P<0.01, ***P<0.001 ****P<0.0001.

### Transduction efficiency was inversely correlated with variant yield

Several of the capsid chimeras routinely produced lower than wild-type AAV6 viral titers from packaging cell lines, while other variants showed no detectible reduction (Fig. 4A). When the T cell transduction performance of each capsid variant was compared to their viral titer, the 3 best performing variants, 2, 4, and 5, were on average 3 logs lower than wild-type AAV6. Comparison of the viral titer of the purified wild-type AAV6 and these three capsid variants by quantitative PCR after DNase I treatment, showed significantly lower titers, 2 to 3 logs, for the three variants (Fig. 4B). Quantification of viral capsids by ELISA, using an antibody recognizing an epitope present on AAV6 and all capsid variants, confirmed that all variants showed a reduced number of viral particles, also at 2 to 3 logs less than wild-type AAV6 (Fig. 4B). This observation indicated that the capsid substitutions in variants 2, 4, and 5 impacted either capsid assembly or the stability of assembled viral particles.

**Figure 4.**
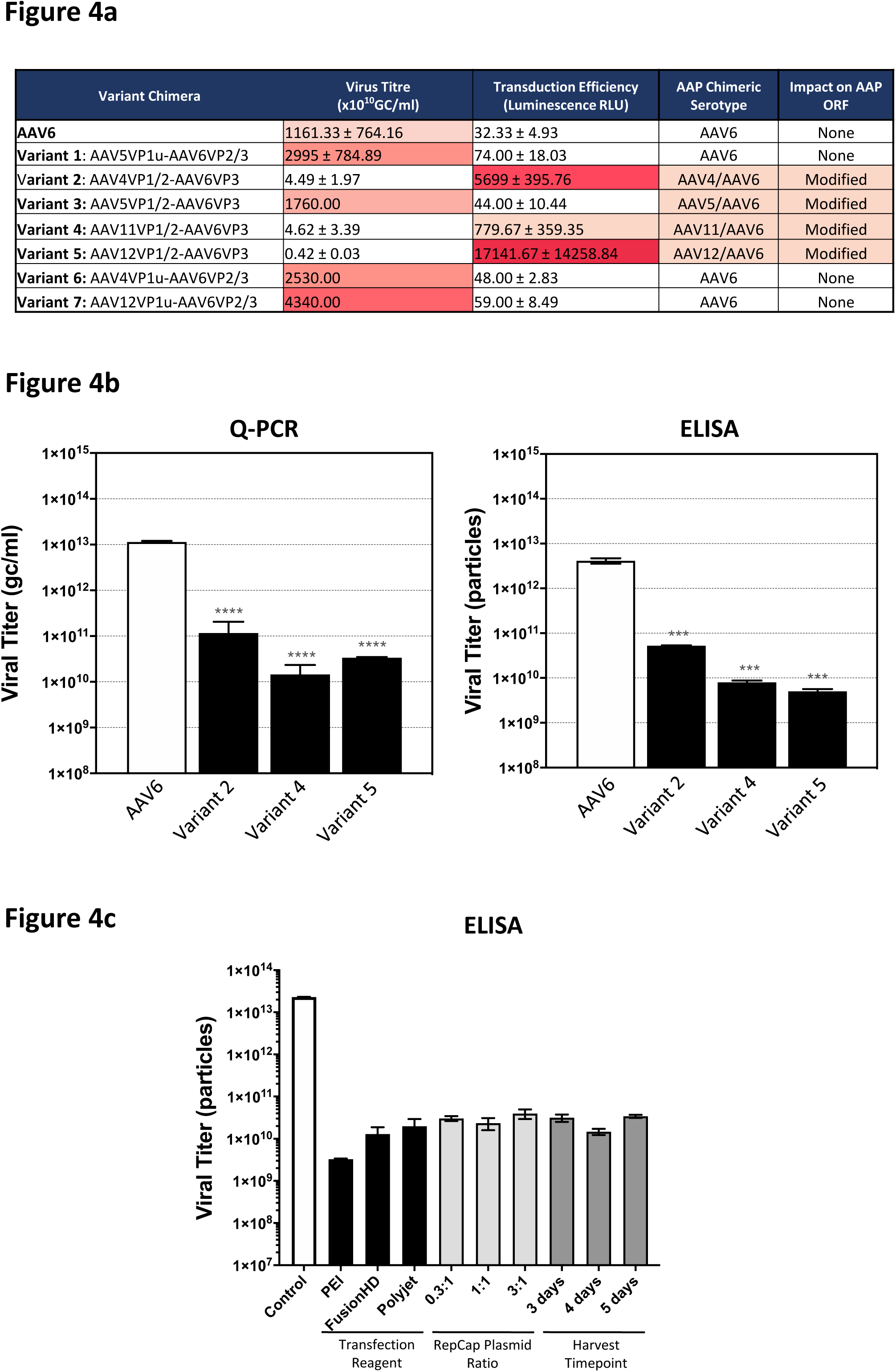
AAV6 capsid variants are defective for capsid assembly. (A) Viral packaging titers and T cell luciferase transduction values for AAV6 capsid variants and the impact of the chimeric capsid sequences on the coding protein sequence of AAP. High rate of transduction correlates with low titer. A heat map coloring is used to indicate the magnitude of yield and transduction. Viral preparations for AAV6 and variants 2, 4, and 5 were produced 3 times; variant 1, 2 times and variants 3, 6, and 7 were produced once. (B) Viral packaging titers from purified AAV particle yield measured by qPCR and ELISA. Both are reduced for variants 2, 4, and 5 compared to wild-type AAV6. (C) Modifying transfection conditions has no effect on AAV6 variant capsid yield. Varying the transfection reagent, the ratio of AAV helper plasmid transfected or the virus harvest time post-transfection, did not lead to any increase in viral genome or particle titer (as measured above) for variant 5. Wild-type AAV6 and variant 5 genome and particle titers were measured from purified particles. Statistical significance was determined by an unpaired T test, or one-way ANOVA test for multiple comparisons of results in triplicate or duplicate replicas, ***P<0.001, ****P<0.0001. GC, Genome copies; RLU, Relative light units.

Attempts to improve variant 5 capsid yield by either optimizing AAV vector transfection method, varying the ratio of AAV helper and AAV donor plasmid transfected, or by extending duration of cell transfection prior to harvesting from the supernatant, failed to demonstrate any detectible increase in viral particle titer (Fig. 4C). All conditions tested resulted in similar vector yield as measured by qPCR (not shown) and average purified particle yield as measured by ELISA (Fig. 4C). Variant 5 capsid yield remained at ~2 to 3 logs lower than for wild-type AAV6. Toward understanding the mechanistic reason for this impairment in particle assembly, the region of the AAV6 capsid sequence altered in these capsid chimera variants were investigated.

### The AAP sequence has an impact on the transduction efficiency of AAV6 capsid chimeras

Given the importance of AAP for AAV capsid assembly (24, 25), the alterations in the sequences of each of the capsid variants within their cap ORF coding for AAP was analyzed. In addition to the VP sequence, the AAP sequence was changed relative to wild-type AAV6 in 4 of the 7 capsid chimeras containing the VP1u+VP1/2 common region (Fig. 5A, and data not shown), including the 3 variants showing enhanced transduction at low MOIs (Fig. 1A and 4A). Previous studies identified the importance of the AAP N-terminus in capsid stability and assembly (26, 27). Thus, to restore a potentially lost AAP function for variant 5, a series of substitution mutants were generated in which the AAP-12 residues equivalent to AAP-6 positions aa13 to aa27 were gradually reverted back to AAV6 (Fig. 5A). A complete rescue of viral titer, to levels of wild-type AAV6, was achieved by reintroducing AAP-6 (Variant 5.1) (Fig. 5B). This observation confirmed the hypothesis that variant 5 has an assembly defect due to compromised AAP function. Approximately 1 log rescue was observed in vector production, compared to variant 5 (AAV-XV), when AAP-12 sequences corresponding to AAP-6 residues aa21 to aa27, the start of the AAP-6 hydrophobic region, were reintroduced to the chimera (Variant 5.3). The remaining substitution variants could not effectively rescue the assembly defect (Fig. 5B).

**Figure 5.**
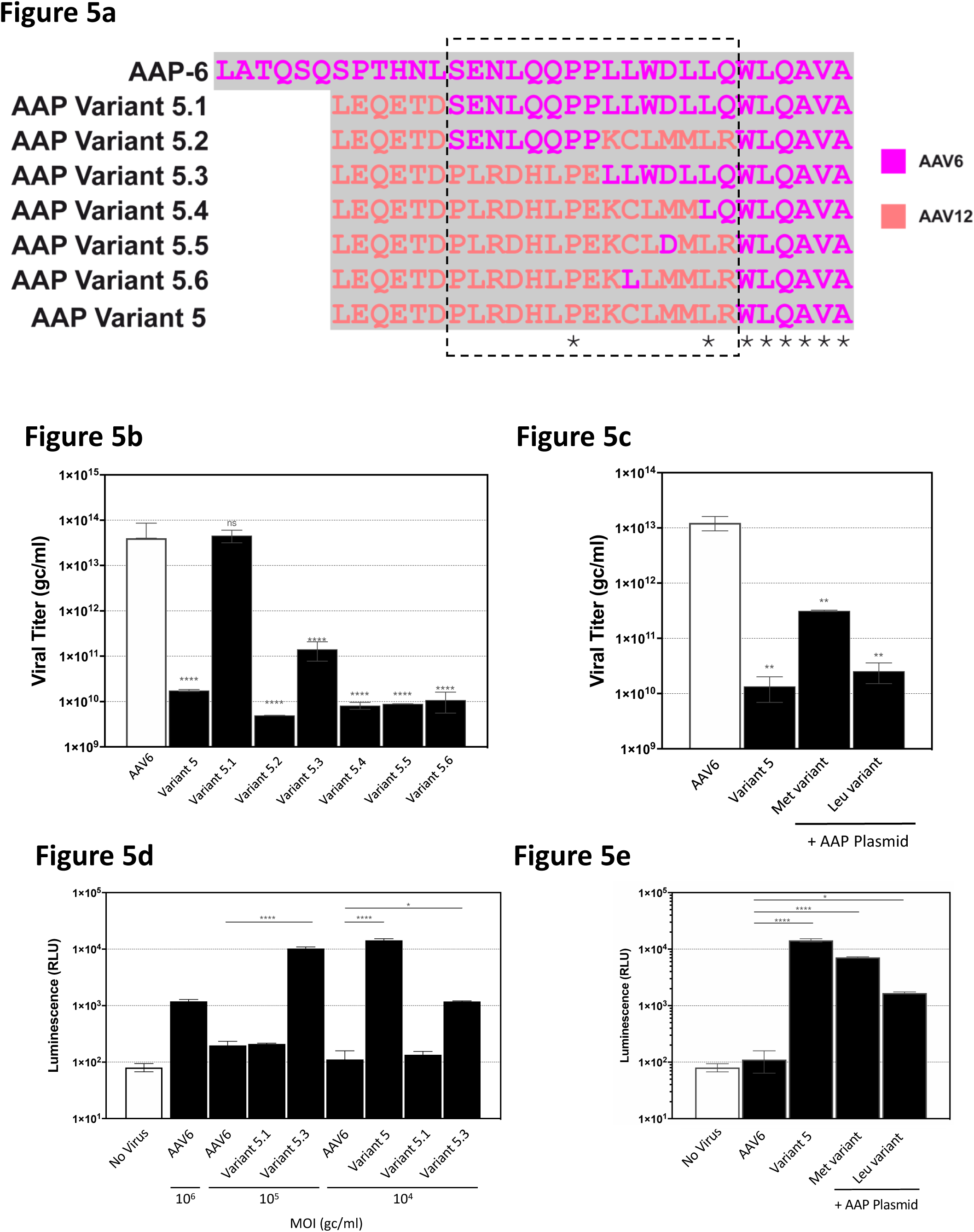
The AAP sequence plays a role in gene expression in addition to capsid assembly. (A) Amino acid sequence alignment of the AAP sequences at the VP2/VP3 boundary for wild-type AAV6 and capsid variant 5 modifications. The wild-type AAV6 sequence is at the top, capsid variant 5 AAP with changes reverting to the wild-type sequence at amino acids 13-27 (outlined by the dashed-line box) of AAP, are 5.1 to 5.6. (B) Genome titers for wild-type AAV6, capsid variant 5, and 5.1 to 5.6 with altered AAP sequences. (C) Genome titer of wild-type AAV6, capsid variant 5 alone and with co-transfection of two forms of the full-length AAP construct in the packaging cells, one starting with a Methionine and the other a Leucine. (D) AAP sequence alterations play a role on transduction of primary CD8+ T cells. Viruses are as in (C). Complete correction of amino acids 13-27, as in variant 5.1, decreases T cell transduction levels, whereas partially corrected 5.3 along with the original variant 5 surpass wild-type AAV6 at low MOI. (E) T cell transduction by variant 5 packaged by co-transfection of the AAP expression constructs (as in D). Statistical significance was determined by an unpaired T test, or one-way ANOVA test for multiple comparisons of results in triplicate or duplicate replicas, *P<0.05, **P<0.01, ***P<0.001 ****P<0.0001.

We further demonstrated that co-transfection with a functional full-length CMV promoter-driven AAP-6 gene into the producer cells, in *trans*, along with the AAV-XV Rep/Cap and donor vector, rescued titer by up to 1.5 logs (Fig. 5C). The AAP gene with the natural CTG start codon (leucine) present in the AAP-6 sequence was insufficient to provide this rescue, while substitution to an ATG start codon to code a methionine, produced the titer rescue (Fig. 5C). This likely reflects the requirement to have a canonical start signal for robust translation initiation and expression of AAP when present within an expression plasmid in HEK293 packaging cells.

Finally, the AAV12VP1/2-AAV6VP3 variant that was fully (variant 5.1) or partially (variant 5.3) rescued by AAP-6 sequence restoration were assessed for T-cell transduction and genomic integration of the luciferase gene when co-delivered with CRISPR (Fig. 5D). Variant 5.1, in which AAP-6 residues aa13 to aa27 had been reverted to AAV6, gave the same level of transduction as AAV6 at all MOIs compared, and had thus lost any enhancement in infectivity or transduction (Fig. 5D). In contrast, variant 5.3, in which only aa21 to aa27 of AAP-6 were reintroduced, maintained some level of superiority over AAV6, resulting in 2 log higher transduction at MOI of 1×10^5^ gc/cell and 1 log higher transduction at the lowest MOI tested of 1×10^4^ gc/cell. Variant 5 maintained superiority to AAV6 at MOI 1×10^4^ gc/ml (Fig. 5D).

Variant 5 vectors, generated with the Met-AAP-6 co-transfection construct, resulted in levels of T cell transduction approximately 2 logs higher than AAV6, and is almost as high as the level observed for variant 5 that is AAP-disrupted (Fig. 5E). The Leu-AAP-6 co-transfection produced sample shows transduction that is ~1 log higher than wild-type AAV6 (Fig. 5E). Collectively these data demonstrate that a modified AAP sequence in the capsid chimera variants can restore vector production at the expense of transduction efficiency at lower MOIs. However, co-transfection with AAP in *trans* during vector production also partially restores packaging (>1 log) and with minimal impact on transduction at low MOI (Fig. 5C and E).

## DISCUSSION

Recombinant AAV vectors hold great promise as a gene delivery vehicle (1, 2). Despite demonstrated clinical efficacy reported for several indications, limitations remain that impede the broader applicability of the AAV technology for efficient and persistent gene delivery to many cell types. The low frequency genomic integration of AAV had previously made the genetic engineering of cells a difficult and laborious task that required the use of drug selection genes or fluorescent markers to select the small number of cells successfully modified with the AAV donor (28). The advent of designer nuclease technologies and the combination of CRISPR gene editing with AAV, enabled substantially higher levels of targeted genome integration of AAV donors and opened up the ability to genetically modify cell types such as primary human T cells for therapeutic applications (17, 18). However, even with these advances in technology, high MOI doses of AAV are still required to reach sufficient levels of transgene integration. Despite optimized production procedures, high dose requirements of AAV are expensive and limit scaling to clinical manufacture. Perhaps even more importantly, recent evidence of immune responses directed towards large quantities of AAV administered *in vivo* to patients in gene therapies highlights the potential toxicity issues associated with high AAV doses (3, 4). In an effort to expand and improve the transduction efficiency and genetic engineering potential of AAV, our chimeric AAV variant design approach, combining VPs from several divergent serotypes, has generated an AAV, AAV-XV, with enhanced transduction characteristics for human T cells that address the requirement for high MOI. AAV6 is considered the best serotype for the transduction of T cells, and thus served as a starting point to evaluate the impact of VP1 and VP2 swapping (Fig. 1) in an attempt to identify modifications that further improved this T cell tropism (19).

Collectively, our data show that an engineered capsid chimera, AAV-XV, comprising AAV12VP1/2-AAV6VP3, can transduce T cells at doses 2 logs lower MOI than wild-type AAV6 with equal efficiency. Both as a donor template in combination with CRISPR gene editing, and as a method for AAV-mediated homologous recombination in the absence of targeted genomic cleavage, superior transduction and stable gene delivery was observed for AAV-XV when compared to AAV6 (Fig. 2 and 3). While the mechanism for this enhanced transduction is not yet clear, the enhanced transduction efficiency requires the C-terminal amino acid sequences of the AAV12 VP1/2 common region. The VP1/2 common region as well as the N-terminus of VP3 are believed to be located in the interior of the capsid and becomes externalized to the capsid surface upon acidification of the endosome during cellular trafficking (29). Thus, the observed enhanced efficiency of infection is likely to be a post-entry effect due to improved interaction with trafficking receptors/effectors. Furthermore, this region has been described as structurally highly flexible (30) and no significant structural differences are predicted (http://original.disprot.org/) by substituting AAV6 to AAV12 sequences.

What is clear from the data is that the modification of the AAV6 capsid sequence to incorporate the VP1/2 of AAV12, results in a change of the amino acid sequence for the overlapping AAP. This is likely the reason for the lower titer yield for the AAV vectors with the variants compared to the wild-type AAV6 given the well characterized assembly-promoting activity of AAP and identification of mutations within this sequence that reduces the interaction of AAP with the capsid, impairing its ability to promoter particle assembly (24–26, 31, 32). The critical region affected in the AAV-XV variant lies within a hydrophobic region of AAP and aa13-27 appear particularly important for stability and assembly functions (26). The data demonstrates that the defect can also be partially rescued by restoring aa21-27 to AAP-6, supporting the theory that these capsid changes impaired AAP-mediated capsid assembly.

The overlapping ORF encoding the VPs and AAP add complexity to the rational design of capsid variants (24). This was evident in the chimeras generated in this study in which VP changes resulted in AAP modification, and low vector yield. However, the ability to transduce cultured human T cells at several log orders lower MOIs to achieve efficient genomic integration of AAV donor DNA requires far less virus to be manufactured, achieving a balance between potency and yield. In conclusion, our data has shown that AAV capsid chimeras generated by VP protein combinations from divergent serotypes is an effective approach for generating novel AAV variants with unique and enhanced functional properties for cellular transduction. Careful consideration of the precise sequence changes is important given the overlapping nature of AAV ORFs and particular care is needed to avoid detrimental modifications to the AAP or MAAP protein (33). The AAV-XV capsid chimera, AAP rescued or not, shows the useful property of highly efficient transduction of cultured human cells at 100-fold lower MOIs compared to the parental AAV6. AAV-XV thus has valuable properties for low-dose gene delivery, enhancing the safety profile of AAV vectors administered *in vivo* to avoid the toxicity that can occur with current AAV-based therapies at high dose.

## METHODS

### AAV variant design and plasmid generation

AAV variants were designed by first extracting the sequences of the VP1u and VP1/2 common region from AAV1 through AAV13 (included AAVrh.10) and performing a pairwise alignment of each serotype to AAV6 using Clustal Omega (https://www.ebi.ac.uk/Tools/msa/clustalo/). The pairwise identity and similarity for each serotype sequence was compared and serotypes with the lowest identities within the VP1u and VP1/2 common region, (serotypes AAV4, AAV5, AAV11, and AAV12) were used to generate the chimeric sequences with the AAV6 VP3. At this point these chimeras were called variant 1 to 7. DNA encoding the chimera regions were generated by DNA synthesis (GenScript Biotech) and subcloned into the AAV6 RepCap plasmid to replace the AAV6 sequence (Plasmid Factory).

A gene targeting vector to measure genome integration via AAV homologous recombination, was constructed by flanking the NanoLuc luciferase gene under the control of the CMV promoter, with 1kb sequences homologous to a transcriptionally active intronic region within the human AAVS1 locus. This AAVS1 luciferase donor contained AAV2 ITRs for packaging of the single-stranded vector into chimeric AAV6 particles, and the ampicillin resistance gene.

### AAV production, purification, and quantification of the genomic titer

Recombinant AAV6 variants were produced by ViGene Biosciences by triple transfection of adherent growing HEK293 cells with the AAV6 variant RepCap plasmids, helper plasmid, and NanoLuc luciferase donor plasmid using polyethylenimine. The transfected cells were harvested 72 h post transfection, pelleted, and subjected to three freeze-thaw cycles to release the AAV vectors from the cells. Vectors released into the grows medium during the 72 h incubation period were recovered by addition of polyethylene glycol (PEG) to a final concentration of 8.2% (w/v) and subsequent precipitation. The rAAVs from the cell pellet and the PEG precipitate were combined and treated with benzonase for 30 min to 2 h. The raw lysate was clarified by centrifugation and the supernatant purified by iodixanol gradient ultracentrifugation, as previously described (34), using a Beckman VTI 50 rotor at 48,000 rpm for 2 h. The genome-containing capsids were extracted from the 40% iodixanol fraction which buffer-exchanged and concentrated using an Amicon^®^ Ultracel 100 kDa cut-off concentrator column (Millipore).

The packaged genome titers were determined by quantitative PCR (qPCR) using Sybr Green stain, with primers directed to the AAV2 inverted terminal repeat regions. The physical particle titer was determined by an AAV6 titration ELISA (Progen, PRAAV6) according to the manufacturer’s instructions, that recognizes a conformational epitope present on AAV6 and all other capsid variants tested here, but does not detect unassembled capsid proteins.

### T cell transduction and luciferase assay

. Primary human CD3+ and CD8+ T cells were isolated from unfractionated PBMCs using the EasySep Human T cell Isolation Kit and Human CD8 T cell isolation kit with RapidSpheres (Stemcell Technologies). Mobilized human primary CD34+ cells from Peripheral Blood (were obtained from Caltag Medsystems, Buckingham, UK). Both T cells and CD34+ HSCs were cultured in X-Vivo 15 media (Lonza) supplemented with 10% human serum AB (Merck Sigma-Aldrich), 300 IU/ml IL-2, and 5 ng/mL IL-7 and IL-15 (Peprotech) at 37°C and 5% CO_2_.

For CRISPR + AAV treatments, 2×10^5^ T cells were first stimulated using anti-CD3/CD28 dynabeads (Invitrogen) in complete T cell media for 48 h prior to electroporation. T cells were electroporated with 15 ug Cas9 mRNA (TriLink) and 10 ug AAVS1 specific sgRNA using the Neon electroporator (3×10^5^ in 10ul Neon tip) and pulse conditions 1400 V, 10 ms, 3 pulses. Electroporated T cells were recovered in T cell media for 2 h before addition of purified AAV vectors to the media at MOI ranging from 200 to 1×10^6^ viral particles. The volume of each virus sample was adjusted by diluting in PBS, so each treatment received equivalent volumes, compensating for the lower concentrated vector samples. Media was replaced after 24 h with fresh complete T cell media, and again every 2 to 3 d. For quantification of T cell transduction, luciferase activity of the transduced T cells was measured at 7, 14, and 21 d post-infection. T cells were harvested, and firefly luciferase analyzed using the Dual-Glo Luciferase assay kit (Promega) according to the manufacturer’s instructions. Luminescence was measured using a PHERAstar microplate reader (BMG Labtech).

### Investigation the role of AAP on AAV6 variant production

For experiments investigating the impact of co-expressing wildtype AAP during viral packaging, AAV production protocols were modified to include polyethylenimine co-transfection of an AAP-6 expression plasmid (ORF under the control of the CMV promoter, synthesized by GenScript Biotech), the variant 5 AAV cap plasmid, and an adenoviral helper plasmid into HEK293 in equivalent amounts, along with the AAVS1-Nano Luciferase targeting vector. Constructs expressing the AAP gene with either the native Leucine start codon, or a substituted Methionine start codon were tested. Virus produced using expression of AAP *in trans* were purified as described above.

For experiments investigating the reversion back to wild-type AAP-6 within the AAP modified in the variants, additional vectors were designed and synthesized with combinations of amino acid substitutions with residues 13-27 of the AAP-6 sequence. Virus preparations packaging the AAVS1-Nano Luciferase construct using each AAP-modified variant were generated and evaluated for transduction in human T cells as described above.

## Supplementary

**S1**. Amino acid capsid sequence alignment of the AAV6 chimera variants with wild-type AAV6. Below a table of the amino acid sequence identities of AAV4, −5, −11, −12 and the variants to the AAV6 VP1, VP1u, and VP1/2 common region is given, respectively.

