## Supplemental Figure S1 for "AAV Capsid Chimeras with Enhanced Infectivity reveal a core element in the AAV Genome critical for both Cell Transduction and Capsid Assembly"

### Supplementary Figure S1

```
        VP1
        1       10        20        30        40        50        60        70
     AAV6 MAADGYLPDWLEDNLSEGIREWWDLKPGAPKPKANQQKQDDGRGLVLPGYKYLGPFNGLDKGEPVNAADAAA
Variant 1 MSFVDHPPDWLEE-VGEGLREFLGLEAGPPKPKPNQQHQDQARGLVLPGYNYLGPGNGLDRGEPVNRADEVA
Variant 2  MTDGYLPDWLEDNLSEGVREWWALQPGAPKPKANQQHQDNARGLVLPGYKYLGPGNGLDKGEPVNAADAAA
Variant 3 MSFVDHPPDWLEE-VGEGLREFLGLEAGPPKPKPNQQHQDQARGLVLPGYNYLGPGNGLDRGEPVNRADEVA
Variant 4 MAADGYLPDWLEDNLSEGIREWWDLKPGAPKPKANQQKQDDGRGLVLPGYKYLGPFNGLDKGEPVNAADAAA
Variant 5 MAADGYLPDWLEDNLSEGIREWWALKPGAPQPKANQQHQDNGRGLVLPGYKYLGPFNGLDKGEPVNEADAAA
Variant 6  MTDGYLPDWLEDNLSEGVREWWALQPGAPKPKANQQHQDNARGLVLPGYKYLGPGNGLDKGEPVNAADAAA
Variant 7 MAADGYLPDWLEDNLSEGIREWWALKPGAPQPKANQQHQDNGRGLVLPGYKYLGPFNGLDKGEPVNEADAAA

                                                                       VP2
              80        90        100       110       120       130       140
     AAV6 LEHDKAYDQQLKAGDNPYLRYNHADAEFQERLQEDTSFGGNLGRAVFQAKKRVLEPFGLVEEGAKTAPGKKR
Variant 1 REHDISYNEQLEAGDNPYLKYNHADAEFQEKLADDTSFGGNLGKAVFQAKKRVLEPFGLVEEGAKTAPGKKR
Variant 2 LEHDKAYDQQLKAGDNPYLKYNHADAEFQQRLQGDTSFGGNLGRAVFQAKKRVLEPLGLVEQAGETAPGKKR
Variant 3 REHDISYNEQLEAGDNPYLKYNHADAEFQEKLADDTSFGGNLGKAVFQAKKRVLEPFGLVEEGAKTAPTGKR
Variant 4 LEHDKAYDQQLKAGDNPYLRYNHADAEFQERLQEDTSFGGNLGRAVFQAKKRVLEPLGLVEEGAKTAPGKKR
Variant 5 LEHDKAYDKQLEQGDNPYLKYNHADAEFQQRLATDTSFGGNLGRAVFQAKKRILEPLGLVEEGVKTAPGKKR
Variant 6 LEHDKAYDQQLKAGDNPYLKYNHADAEFQQRLQGDTSFGGNLGRAVFQAKKRVLEPLGLVEQAGETAPGKKR
Variant 7 LEHDKAYDKQLEQGDNPYLKYNHADAEFQQRLATDTSFGGNLGRAVFQAKKRILEPLGLVEEGVKTAPGKKR

                                                                        VP3
             150        160             170       180       190       200
     AAV6 PVEQSPQE---PDSSSGIGKTGQ-----QPAKKRLNFGQTGDSESVPDPQPLGEPPATPAAVGPTTMASGGG
Variant 1 PVEQSPQE---PDSSSGIGKTGQ-----QPAKKRLNFGQTGDSESVPDPQPLGEPPATPAAVGPTTMASGGG
Variant 2 PLIESPQQ---PDSSTGIGKKGK-----QPAKKKLVFEDETGAG---DGPPEGSTSGA--------MASGGG
Variant 3 IDDHFP-----KRKKARTEEDSK--------PSTSSDAEAGPSGS----QQLQIPAQPASSLGADTMASGGG
Variant 4 PLES-PQE---PDSSSGIGKKGK-----QPARKRLNFEEDTGAG---DGPPEGSDTSA--------MASGGG
Variant 5 PLEKTPNRPTNPDSGKAPAKKKQKDGEPADSARRTLDFEDSGAG---DGPPEGSSSGE--------MASGGG
Variant 6 PVEQSPQE---PDSSSGIGKTGQ-----QPAKKRLNFGQTGDSESVPDPQPLGEPPATPAAVGPTTMASGGG
Variant 7 PVEQSPQE---PDSSSGIGKTGQ-----QPAKKRLNFGQTGDSESVPDPQPLGEPPATPAAVGPTTMASGGG
```

| | amino acid sequence identity vs. AAV6 (in %) | | |
|---|---|---|---|
| | VP1 | VP1u | VP1/2 |
| AAV4 | 63 | 87 | 46 |
| AAV5 | 58 | 72 | 22 |
| AAV11 | 65 | 99 | 46 |
| AAV12 | 60 | 89 | 30 |
| Variant 1 | 95 | 72 | 100 |
| Variant 2 | 93 | 87 | 46 |
| Variant 3 | 88 | 72 | 22 |
| Variant 4 | 96 | 99 | 46 |
| Variant 5 | 91 | 89 | 30 |
| Variant 6 | 98 | 87 | 100 |
| Variant 7 | 98 | 89 | 100 |
